# Influence of human population density on spatial distribution patterns of environmental suitability for triatomine vectors of Chagas disease

**DOI:** 10.1101/717348

**Authors:** Anderson A. Eduardo, Lucas A. B. O. Santos, Mônica C. Rebouças, Pablo A. Martinez

**Author notes:** Corresponding author. E-mail (AAE).

## Abstract

Previous work on Chagas Disease disease at large spatial scales has not explored how interaction with humans can affect projections for geographical distribution of environmental suitability of vector species. Here, we compare niche-based species distribution models with climatic variables as predictors (SDM_clim_) and with climatic variables + human population density (SDM_Human_). Our results show that accounting for human population density helps refine the models to finer geographical scales. Also, different spatial patterns of accumulated environmental suitability were obtained by SDM_clim_ and SDM_Human_. Moreover, projections were more accurate for SDM_Human_ than for SDM_clim_. Our results show that considering human populations in SDMs for epidemiologically relevant triatomiane species can improve our understanding of macroecology and biogeography of environmental suitability for vectors of Chagas disease.

## 1. Introduction

Infectious diseases cause approximately 9.6 million deaths each year, resulting in economic costs which can reach the order of billions of dollars/year and invaluable social costs [1–4]. From this universe, a group of etiological agents that have been little studied makes a disproportionate contribution to the global burden of human disease, being reported by the World Health Organization (WHO) as neglected tropical diseases (NTDs) [3,5]. They are among the major diseases affecting poor populations around the world, being estimated that about 1 billion people are infected by one of the 17 NTDs [2,5,6]. In Latin America, Chagas disease or American Trypanosomiasis is the main NTD in most countries, presenting the greatest social impacts among all infectious diseases [1,2,5]. In Brazil, the South American country with the greatest number of NTDs, Chagas disease is the one with the highest case fatality rate [1,2,5–7]. The human infections can occur by blood transfusion and by ingestion of food infected with the etiologic agent (the protozoan parasite *Trypanosoma cruzi*), however the primary form of transmission occurs by the bite of triatomine hematophagous insect, popularly called “kissing bugs” (Hemiptera, Reduviidae) [7]. Due to the lack of vaccine and efficient chemotherapeutic treatments, epidemiological control strategies are focused on the control of vector infestations [8,9]. Despite the relative success of these actions, *T. cruzi* infection through triatomine vectors continues to represent a critical transmission route for the epidemiology of Chagas disease, with occurrences of home and peridomiciliary infestations, as well as records of new acute cases each year [10]. In this context, research that contributes to increase the consistency of the theoretical background of vector control practices remains needed.

Most ecological studies on the vectors of Chagas disease focus on small-scale, environmental factors affecting species distribution and abundance locally (i.e., household, peridomestic and wild microenvironments) [11–16]. Only in recent years, research inquiring about the large-scale spatial patterns of species distribution and species richness of triatominae vectors become available in the literature [10.17–23]. These works contribute importantly not only to our understanding of Chagas disease macroecology but also to remedy the generalized gap of knowledge about the determinants of infectious diseases at the macroscales of biological organization [3,4,24]. However, until now these studies focused on the influence of major climatic factors and anthropogenic climatic changes on geographical distribution of triatomine vectors of Chagas disease. As vectors of an infectious disease, the ecological interaction among triatomine species and humans is conceptually inherent and the evaluation of how much this interaction can influence environmental suitability for triatomine occurrence presents a relevant task in disease ecology and spatial epidemiology.

Most of the research regarding triatomine species at macroecological and biogeographical scales rely on the use of niche-based species distribution models (SDMs). Essentially, SDMs correlate species occurrence data with environmental variables across the geographic space of their distribution. This information is used by computational algorithms able to elaborate a model of the ecological niche of the focal species and to make predictions in the geographic space [25–27]. Implicitly, SDMs assume that the species is in equilibrium with the environment, i.e., the species currently occupies all those areas suitable for it [25,27], and that climatic niches are conserved through recent geological time [25,27]. Despite the wide range of applications, models have been consistently used to capture the species’ Grinellian niche (or climatic niche) [28]. Recently, some authors have invested efforts to appropriately include biotic interactions in SDMs [29,30]. This is a promising methodological approach and research on vectors’ ecology can directly benefit from it. Species interactions represents an important aspect for the spatial distribution of any infectious disease vector, since work over the last few years (e.g., [31–34]) have demonstrated that biotic interactions (including with the human species) can be influential at macroecological scales, which can lead to erroneous predictions if not properly considered in SDM implementation. Moreover, such biotic interactions should be of particular of interest for the predictive accuracy of environmental suitability projections for other moments of time, such as future climatic scenarios for example [34,35].

Previous studies has shown that triatomines can have different levels of adaptation to anthropogenic environments [13,22]. They are organisms that can move around actively looking for microenvironments that meet their ecological demands and, thus, the species can infest best suited household and peridomestic areas [22,23]. These different levels of successful colonization of anthropic environments can be interpreted as different levels of ecological interaction between different species of triatomine vectors and the human species. Given the prevalence of humans in the Earth’s ecological systems, this interaction may represent a critical aspect of the niche of these species, influencing environmental suitability throughout geographical space and, potentially, affecting the spatial pattern of species distribution.

In this context, in the present study our objective was to investigate the influence of human species on environmental suitability of kissing bugs, triatomine vectors of Chagas disease. Specifically, we focused on epidemiologically relevant species in Brazil and assessed how the inclusion of spatial data of human population density affect the projections of niche-based spatial distribution models (SDMs) based in climate variables. Considering the relevance of Chagas disease in Latin America, the results of such analysis are important to the consistency of the knowledge that underpins the decisions regarding epidemiological control strategies.

## 2. Materials and methods

### 2.1 Occurrence data

Through the platforms Species Link (www.specieslink.org.br), GBIF (Global Biodiversity Information Facility – www.gbif.org), as well as searching directly in the literature, we obtained a total of 609 occurrence records for 14 of the 15 triatominae species reported as epidemiologically relevant by the Brazilian Department of Health Surveillance (*Panstrongylus geniculatus, Panstrongylus lutzi, Panstrongylus megistus, Rhodnius nasutus, Rhodnius neglectus, Rhodnius robustus, Rhodnius pictipes, Triatoma infestans, Triatoma brasiliensis, Triatoma maculata, Triatoma pseudomaculata, Triatoma rubrovaria, Triatoma sordida* e *Triatoma vitticeps*) [8,9]. Considering the findings of other authors (specially Proosdij et al. [36]), we assumed 20 records as the minimal sample size for our analysis. So, *Triatoma rubrofasciata* was not considered in our analysis because we were able to compile less than five records throughout the our study area. Records not identified at taxonomic level of species (e.g., identified at genus taxonomic level), with uncertainty in coordinates greater than 5 km, points of occurrence outside of continental areas, as well as repeated points at the same site (i.e., same pixel in our raster layers, having a resolution of 2.5 minutes – details in the next section), were excluded from our datasets. Our final datasets are provided at supplementary material. All procedures were performed in R software environment, using packages rgibif (https://CRAN.R-project.org/package=rgbif) and raster (https://CRAN.R-project.org/package=raster).

### 2.2. Climatic variables and human population density data

Following other authors [17–21], we have modeled the niche of each one of out 14 triatomine species using species distribution models (SDMs) built upon climatic variables related to temperature (in °C) and precipitation (in mm). So, we used the 19 bioclimatic variables of the World Clim (www.worldclim.org; [37]), with resolution of 2.5 minutes (≈ 5km × 5km in equatorial regions). In order to minimize multicollinearity, we employed a stepwise procedure to exclude highly correlated variables (i.e., *r* > 0.7). Specifically, first we calculated Pearson pairwise correlation among the 19 bioclimatic variables. Then, the employed algorithm find the pair of variables which has the higher correlation and exclude one of then which has greater VIF (variance inflation factor). This procedure is iterated until no highly correlated variables remains. For convenience, we used the functions from usdm package (https://CRAN.R-project.org/package=usdm) in R software environment for our implementation of variable selection. After this procedure, the following variables were selected: Bio 2, Bio 3, Bio 8, Bio 10. Bio 11, Bio 12, Bio 13, Bio 14, Bio 15, Bio 16, Bio 18, Bio 19 (respectively, mean diurnal range, isothermality, mean temperature of wettest quarter, mean temperature of warmest quarter, mean temperature of coldest quarter, annual precipitation, precipitation of wettest month, precipitation of driest month, precipitation seasonality, precipitation of wettest quarter, precipitation of warmest quarter, precipitation of coldest quarter).

Human population density data were obtained from Grided Population of the World, Version 4 (GPWv4) [38], a spatially explicit database of human population (http://sedac.ciesin.columbia.edu/data/collection/gpw-v4). We used the most recent available data (which currently is for the year of 2015), at 2.5 arc-minutes and clipped for South America. All described procedures were performed in R software environment, using packages raster and usdm (https://CRAN.R-project.org/package=usdm).

### 2.3. Models of species distribution

The niche-based species distribution models (SDMs) for each one of our 14 studied triatomine species were implemented with Maxent [39,40], using the package dismo (https://CRAN.R-project.org/package=dismo) in R software environment. To consider the interaction with humans in the SDMs we rely on the framework provided by Anderson [30]. According to this author, variables influencing species’ environmental suitability can be thought as related to two groups: variables which dynamics are unlinked to focal species dynamics (scenopoetic variables, *sensu* Hutchinson [41]) and variables which dynamics are linked to the dynamics of the focal species. In this sense, ecological interactions among a pair of species in which one of them is unaffected by the other are prone to be captured by SDMs in a straightforward fashion. In such cases, when the affected species is the focal one for a SDM implementation, data related to abundance for the unaffected species can be included among the predictor variables in the model. This is exactly the case for the interaction between triatomines and humans, in which human population density can be considered unlinked to the population dynamics of triatomines. Relying on this conceptual framework, we built two types of models for each of the triatomine species: the SDM_clim_, only with climatic variables; and the SDM_Human_, with climatic variables + human population density.

To implement each of them, for each species background points were generated through sampling 10,000 points across South America geographical area using a bias layer [42–44]. This procedure is one of the most recommended frameworks when accounting for spatial bias in the focal species occurrence data [43,44]. Our bias layer were produced using a spatial kernel density estimation algorithm (specifically, we used the function kde2d from the package MASS, in R software environment) and employing data for all Reduviidae occurrences available at GIBIF. Presences and background points were grouped together to form a consolidated dataset (i.e., one dataset for each one of our 14 focal triatomine species). Posteriorly, the set of points in each dataset were splitted in geographical blocks (or bins) with approximately sizes. So, the model can be calibrated using data from three blocks and validated with data from remaining one. This process were iterated until each block were used in the validation step. We opted for this method because it has been pointed out as the most effective for evaluating the performance of SDMs [45,46]. Also, we employed the partial area under the Receiver Operating Characteristic curve (pROC) as a measure of model performance. This is a metric related to AUC (Area Under the ROC curve), which is threshold independent metric of model ability in assigning high suitability to those areas where the species is actually present and lower suitability to those areas where the species is actully absent. However, AUC performs in a biased fashion for SDMs [47,48] and the use of pROC have been advocated as an better approach [45]. In practical terms, pROC allows the modeler to define a biologically acceptable level of omission in the performance of the SDM. In the present work, we use an acceptance limit of 5% of omission error. Here, we used the R function PartialROC from the package ENMGadgets (https://github.com/narayanibarve/ENMGadgets) to compute pROC for each species-specific SDM.

Regarding Maxent parametrization, we assessed model complexity (i.e., combinations of Maxent features and regularization multipliers; see [49,50]) testing over each possible combination of features (being allowed linear, linear + quadratic, hinge, linear + quadratic + hinge, linear + quadratic + hinge + product, linear + quadratic + hinge + product + threshold, respectively) and regularization multipliers (ranging from 0.5 to 5.5, by steps of 0.5). The outputs of each model parametrization were compared through AICc scores and the one with the lowest scores were used. Then, for each species, SDMs with the selected parametrization were used to implement spatial projections of environmental suitability (using all occurrence data) along the geographical area inside an estimated accessible area. Following other authors [51], we hypothesized that the accessible area for one species is that within the minimum convex polygon described by its occurrence points in the geographical area plus a buffer equal to the mean distance between each occurrence point and the centroid of the set of points. Finally, SDMs’ projections were converted in binary maps (0=non-habitat; 1=suitable habitat) using species specific threshold based on omission error [45]. In the present study, we used a omission error level of 5%.

All procedures described here were performed in R software environment using the packages ENMeval (https://CRAN.R-project.org/package=ENMeval), ENMGadgets (https://github.com/narayanibarve/ENMGadgets) and raster.

Given that human population data could be considered as a scenopoetic predictor variable in the SDMs, we infer ecological influence of human density on environmental suitability for each tritaomine species in terms of Maxent metrics of variable importance. Specifically, we rely on Maxent’s permutation importance metric as a measure of the relative influence of each predictor variable for the SDMs. For the computation of permutation importance, Maxent algorithm randomly changes variable values between presence and background points and then measures the impact on model performance. Variables highly influential in model performance will present higher values of permutation importance.

For suitability maps, we measured the difference between SDM_Human_ and SDM_clim_, i.e., SDM_Human_ – SDM_clim_, allowing us to map (species specific) geographical discrepancies in modeled suitability with the inclusion of human density in the models. Here, such maps are filled with values ranging from −1 to 1 (naturally, sites where SDM_Human_ and SDM_clim_ project similar suitability values are maped with values closer to zero). Also, we mapped the overlap between binary maps for SDM_clim_ and SDM_Human_ of each species, inspect the result visually, and calculate the proportion of concordance between maps (which were measured using the overlap area between binary maps produced by SDM_Human_ and SDM_clim_ for each species, as well Pearson correlations for environmental suitability maps produced by each one of this model types for the species). Finally, the accumulated suitability across species (i.e., pixel-by-pixel sum of suitability maps for the modeled species – divided by 14, as for 14 species this is the maximum possible value when summing up suitability from each species; hence our accumulated suitability map can be presented ranging from 0 to 1) were mapped for both kinds of SDMs, allowing us to inspect geographical patterns of environmental suitability for the focal triatomine species. As an additional assessment, we evaluated spatial regressions between records for new cases of Chagas disease and accumulated suitability projections produced by SDM_human_ and by SDMclim. We used data for Brazilian municipalities for the last 10 years (2007-2017), made available by Sistema de Informação de Agravos de Notificação at http://datasus.saude.gov.br/ (see Supporting Material). The software SAM was used for spatial regression [52].

## 3. Results

Our assessment of SDM_clim_ and SDM_Human_ showed that human population density represents an important variable for the models. For most modeled species, SDMs have better performance when human population density is considered among the predictor variables, increasing pROC by 26.68% on average. Moreover, human population density always performed as one of the main variables of the set of the predictor variables used, being the most important one for most of the species (9 of 14 triatomine species). Among this last group of species, the largest differences in model performance (ΔpROC) were observed for *P. geniculatus* and *T. infestans.*

**Table 1:** Performance metrics (pROC) for the implemented species distribution models (SDMs). Species names are arranged in the rows, models and performance metrics are arranged in the columns of the table. SDM_clim_ and SDM_Human_ are the models with and without human population density as a predictor variable, respectively (see Material and Methods section); ΔpROC stands for the difference of pROC between the different models (values in bold represents loss in performance when human population density is added to the SDM); the permutation importance of human population density as a predictor variable in the SDM_Human_ were computed by the Maxent algorithm and are provided in the last column.

| Species | pROC | | ΔpROC | Permutation importance of human pop. density in SDM<sub>Human</sub> |
| --- | --- | --- | --- | --- |
|  | SDM<sub>clim</sub> | SDM<sub>Human</sub> |  |  |
| *Panstrongylus geniculatus* | 0.578 | 0.982 | 0.404 | 90.813 |
| *Panstrongylus lutzi* | 0.836 | 0.987 | 0.151 | 95.536 |
| *Panstrongylus megistus* | 0.686 | 0.979 | 0.293 | 88.034 |
| *Rhodnius nasutus* | 0.792 | 0.993 | 0.201 | 70.068 |
| *Rhodnius neglectus* | 0.759 | 0.992 | 0.233 | 38.106 |
| *Rhodnius pictipes* | 0.736 | 0.863 | 0.127 | 44.538 |
| *Rhodnius robustus* | 0.842 | 0.917 | 0.075 | 27.464 |
| *Triatoma brasiliensis* | 0.764 | 0.99 | 0.226 | 92.154 |
| *Triatoma infestans* | 0.635 | 0.941 | 0.306 | 23.041 |
| *Triatoma maculata* | 0.901 | 0.886 | **-0.015** | 3.900 |
| *Triatoma pseudomaculata* | 0.827 | 0.979 | 0.152 | 90.685 |
| *Triatoma rubrovaria* | 0.822 | 0.904 | 0.082 | 7.065 |
| *Triatoma sordida* | 0.789 | 0.994 | 0.205 | 55.192 |
| *Triatoma vitticeps* | 0.747 | 0.972 | 0.225 | 72.310 |

Geographically, considering the human population density in triatomines SDMs increased to spatially heterogeneous projections of suitability. While SDM_clim_ tended to project suitability more continuously distributed across large geographical areas, SDM_Human_ tended to produce more discontinuous projections throughout the space. This can be verified through the overlap between binary maps (Figure 1; Supplementary Material) and the distribution of suitability difference (Figure 2) between SDM_clim_ and SDM_Human_ of each species.

**Figure 1:**
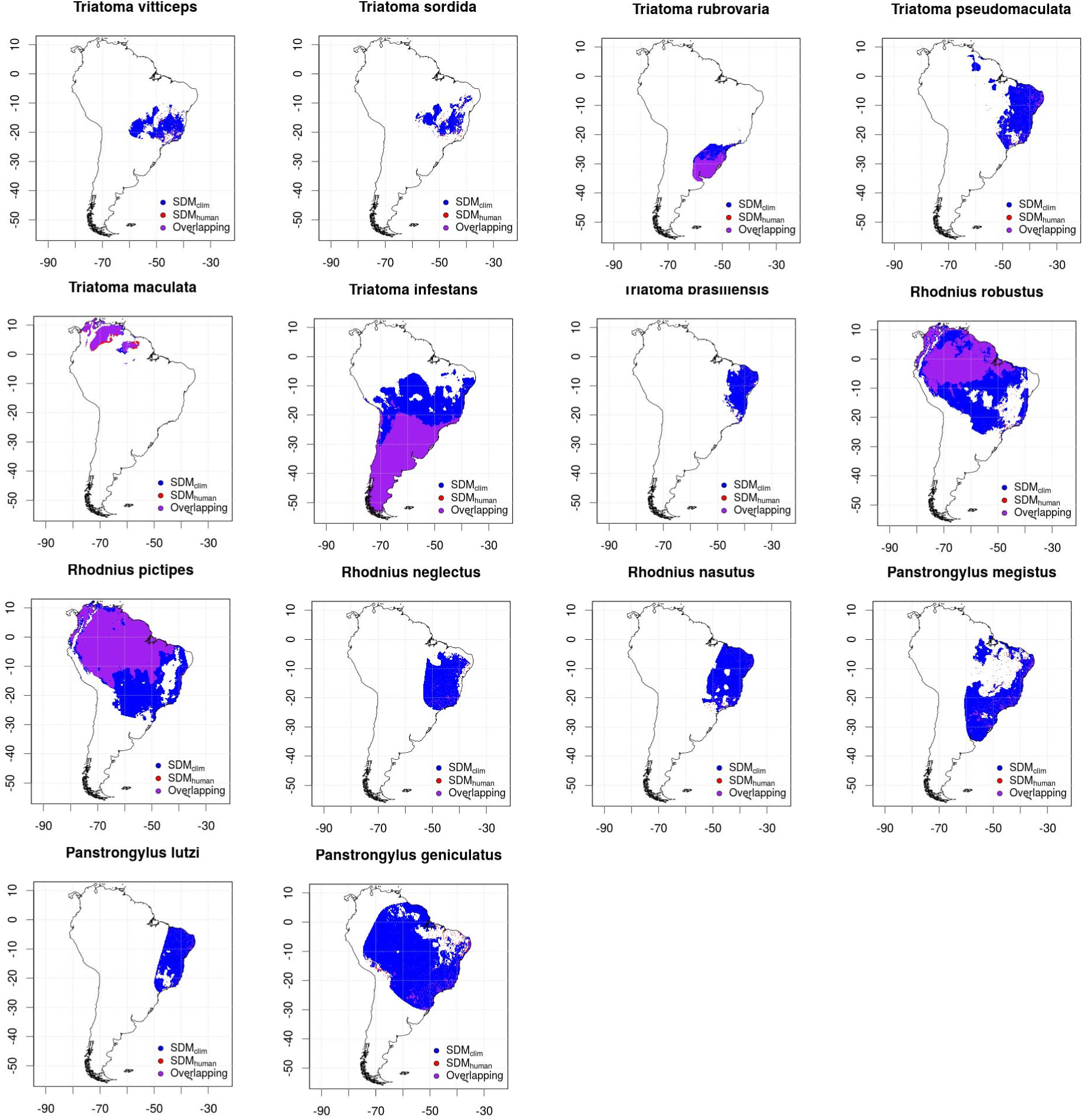
Results of Maxent models for the 14 triatomine species considered in this study. SDM_Human_ and SDM_clim_ are compared through overlapping binary maps of each species.

**Figure 2:**
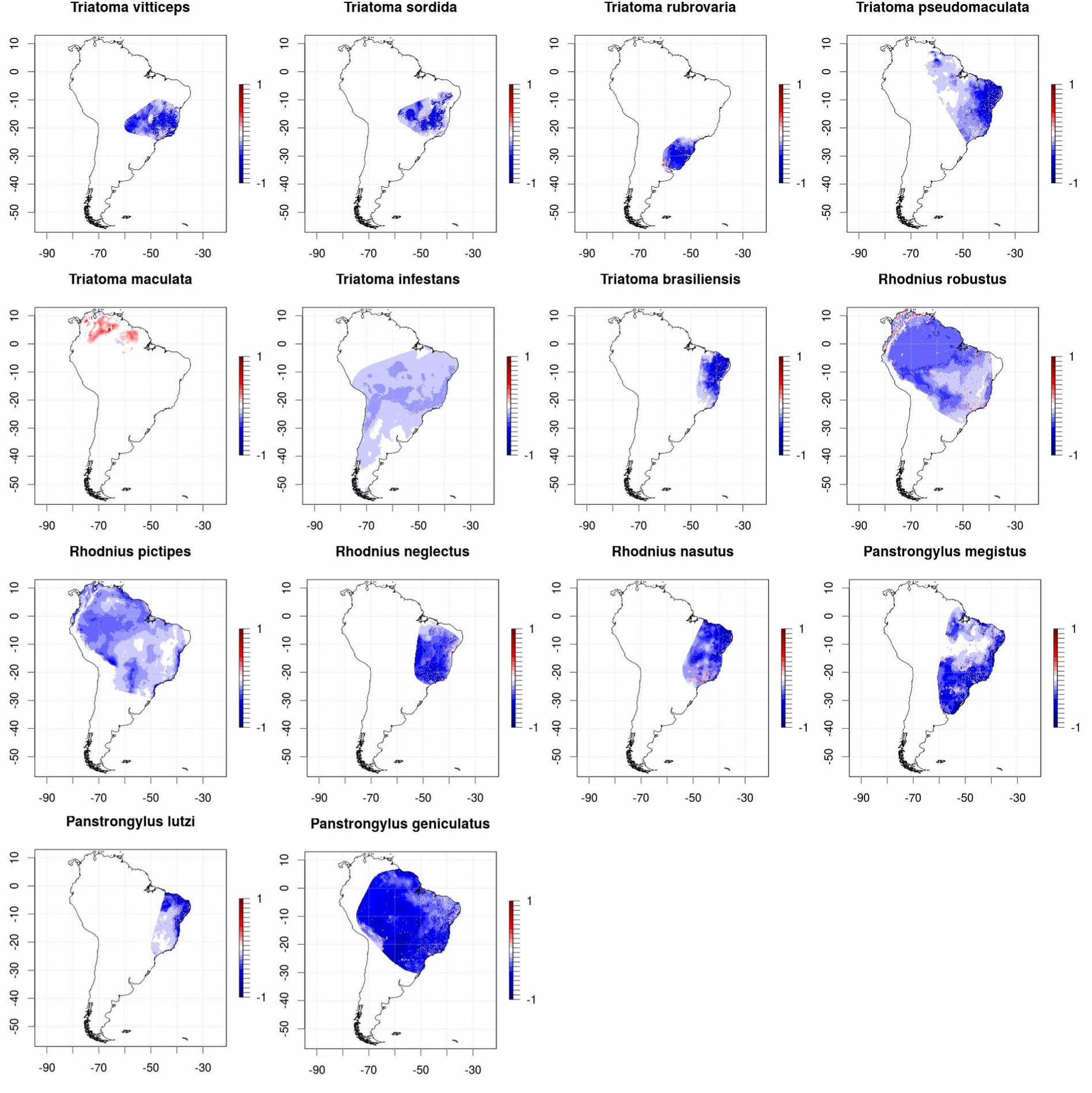
Results obtained from the (pixel-by-pixel) difference between projections of SDM_clim_ and SDM_Human_ for each triatomine species (i.e., suitability map obtained with SDM_Human_ minus suitability map obtained with SDM_clim_). Values in the maps range from −1 (blue) to 1 (red). So, regions where the inclusion of human density in SDMs led to lower predictions of suitability are maped in blue tons. Regions where accounting for human density increases modeled suitability are maped in red tons.

Comparatively, for the species which human population density have higher influence in the models’ performance we observed the lowest overlay in binary maps. For example, for *P. lutzi* and *T. pseudomaculata* we obtained a permutation importance of 91.913 and 88.502 for human population density and binary maps overlap were 0.049 and 0.096, respectively. For *T. rubrovaria* and *T. maculata* we obtained 0.429 and 2.989 for permutation importance and 0.647 and 0.917 for binary maps overlap, respectively. However, in general, this does not represent improvement in SDM performance. The ΔpROC were 0.051, 0.028, 0.008, −0.008 for *P. lutzi, T. pseudomaculata, T. rubrovaria* and *T. maculata*, respectively. Apparently, species with higher delta pROC are the ones with higher differences between SDM_clim_ and SDM_Human_ projections, in terms of geographical extension. However, this is not conclusive.

Beyond the differences in terms of geographical areas, human population density affected environmental suitability distribution (as a continuous response variable) throughout the overlapping areas projected by SDM_clim_ and SDM_Human_. In this sense, we observed that suitability were higher at the southern portion of the modeled distribution of *T. rubrovaria* when human density is considered in the SDM. For *T. maculata*, higher suitability throughout the whole modeled distribution area is obtained with SDM_Human_. For other species with higher overlap (i.e., *R. pictipes, T. infestans, R. robustus*) no clear or relevant patterns were observed. The results for accumulated suitability (Figure 3) shows that high quality habitats for most of the species analyzed are located at northeastern and south-eastern Brazil, despite the consideration of human population density in SDMs. Accounting for human density, we observed that most of the accumulated environmental suitability are associated to localities populated by humans. The accumulated environmental suitability throughout northern and southern Brazil were lower and exhibits a more continuous distribution, not associated to human populated localities. Spatial regression between records of new Chagas disease cases (for Brazilian municipalities, between 2007 and 2017) and accumulated suitability obtained from SDM_Human_ showed a statistically significant positive slope (0.621, *p*-value < 0.05), being positive but not statistically significant (0.622, *p-*value > 0.05) when projections from SDM_clim_ were used. Detailed results are available in Supporting Material.

**Figure 3:**
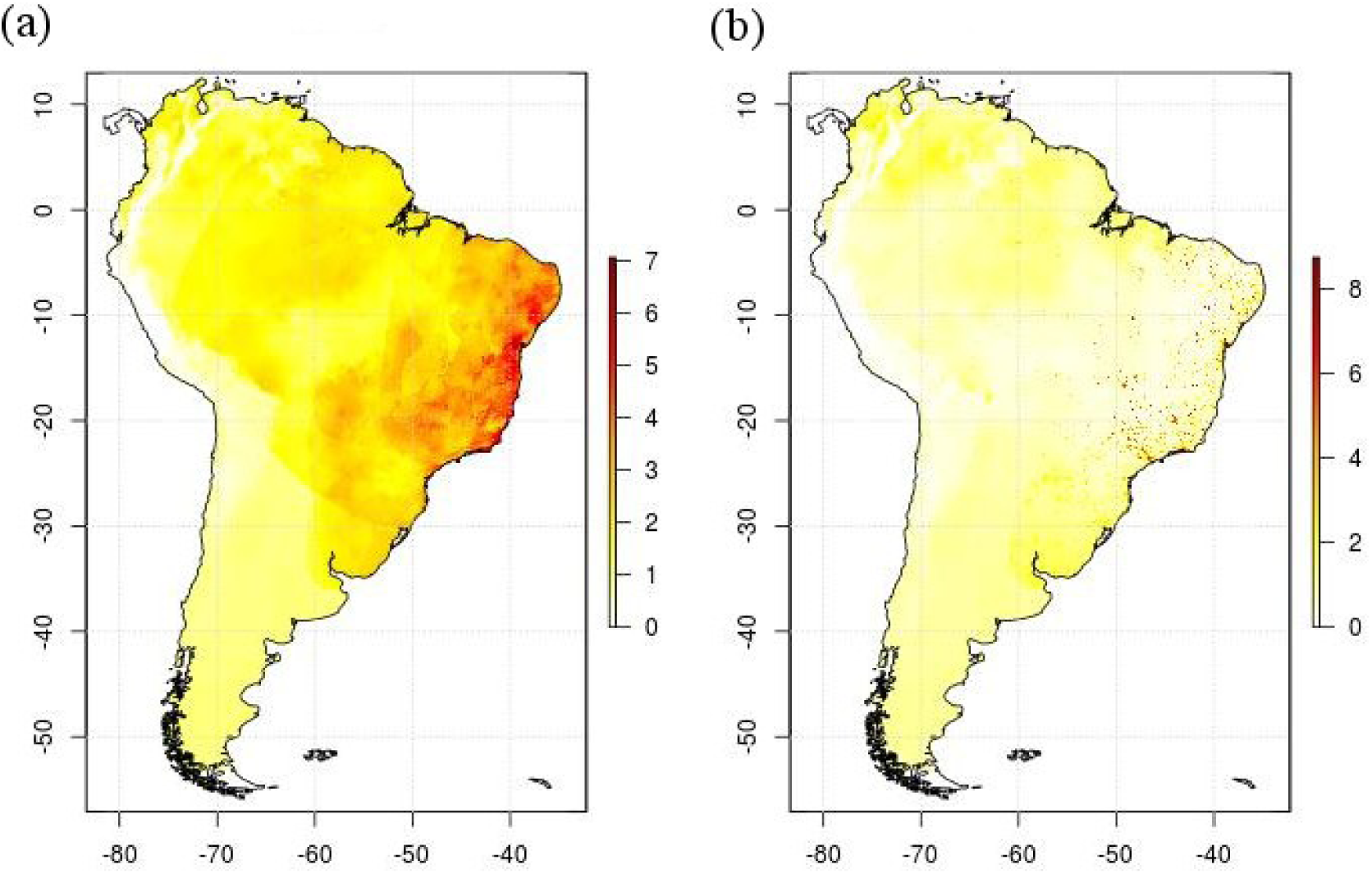
Map of accumulated environmental suitability for epidemiologically relevant triatomine species in Brazil. Each map was produced by adding the suitability map projected for each triatomine species using Maxent. Results for SDM_clim_ (a) and SDM_Human_ (b) are presented. Higher values indicate locations with greater accumulation of suitability (considering the 14 epidemiologically relevant triatomine species studied here) and are indicated in darker shades of red.

## 4. Discussion and conclusions

Together, our results showed that the inclusion of the human population density can affect the projections of environmental suitability obtained from SDMs. Ecologically, human density performs as a relevant environmental covariate for the niche of the analyzed triatomine species. So, on the one hand, human populations trigger a range of environmental alterations which biologically favor (at least) most of the analyzed studied species. These environmental alterations comprehends an amalgam of local conditions, being not it possible here to specify which dimensions of the triatomine species niche were decisive for the differences observed between SDM_clim_ and SDM_Human_. On the other hand, it is precisely the generality of such environmental alteration that allows us to argue that the ecological interaction among human density and triatomine species may represent a relevant aspect for the ecological niche of these vector species. Future work that seeks to thoroughly analyze the nature of anthropogenic environmental changes and how they affect the availability of conditions and resources for triatomine vectors may advance and deepen our results, paving the way for highly detailed projections in time and space to become possible and benefiting the accurate identification of sites with high suitability for proliferation of triatomine vectors.

Despite our focus in triatomine species with epidemiological relevance for Brazil (see [8,9]), the geographical patterns of accumulated environmental suitability mapped by SDM_clim_ is concordant with patterns of species richness provided by other authors who investigated the triatomine group more comprehensively. The most extensive studies were [17] and [20], the first providing an analysis of the species richness patterns of 115 species for the whole Western Hemisphere and the second assessing the thermal tolerance of two species (*T. infestans* and *R. prolixus*), bringing insights into the distribution of the triatomine group as a whole. Specific studies were carried out for Venezuela [19,53], Argentina [53], Colombia [21,23] and Ecuador [54]. For Brazil, in particular, [10] analyzed the distribution of eight species in the center-west region and [55] analyzed the distribution of the species complex *T. brasiliensis*, which occurs mainly in the Brazilian northeast. Such results are in consonance with the species distributions and the patterns of environmental suitability obtained through SDM_clim_.

As evidenced in our results, human population density can affect model projections for geographical distribution of environmental suitability of Chagas disease vectors. Despite of that, even before the advent of large-scale, macroecological studies on triatomine distribution and richness, *T. infestans* was recognized as an emblematic example of the potential of human influence on the geographical distribution of vector species in this group [13]. This is a species with wide distribution in South America, which is pointed out in the literature as a consequence of its high association with humans [13,56]. Due to its ability to completely perform its life cycle and establish persistent populations in anthropic environments, this triatomine species was the main target of measures of epidemiological control of Chagas disease over the last decades [8,9,13]. However, *T. infestans* are not among the species most influenced by human population density in our analysis. In fact, isothermality (i.e., variable Bio 3 variable in our SDMs) was the most influential predictor in the SDMs for this species. We highlight that such apparent discrepancy rely on the different spatial scales involved, which was continental scale in our implemented models whilst classical works on *T. infestans* ecology focused on local scale. This species has historic occurrence records from central Argentina and north Bolivia to the Brazilian Atlantic coast, and the west coast of southern Peru [57,58]. So, niche-based models, focusing on Grinnellian niche, should capture the range of climate conditions in which the species can maintain viable populations [28]. In the case of *T. infestans*, the influence of human species at local spatial scales does not scales up to macro-spatial scales, where climate conditions tend to be critical. This draws attention to the fact that the influence of biotic interactions may not be linearly transposed between different spatial scales during construction and interpretation of environmental suitability models.

It has been widely recognized that the global climate change will have broad impact in the distribution of the infectious diseases [19,59]. The extent to which human populations will suffer the adverse effects of this biogeographical reorganization of diseases depends closely on the consistency of the knowledge that underpins decision makers in public health systems [19,59]. Previous works show that, unlike diseases with direct transmission among humans, zoonoses are not uniformly distributed on the globe, presenting strong geographical bias [60.61]. Furthermore, zoonoses account for most of the emerging infectious diseases in humans [60.61]. In this context, the macroecology and biogeography of species that are related to the epidemiological cycle of etiological agents of infectious diseases assumes a critical aspect in epidemiology. Moreover, it is important to emphasize that both vector species and reservoir species may present different levels of ecological interaction with humans, and accounting for such nuances in SDMs can yield new insights in spatial epidemiology of vector born infectious diseases.

Our results show that more detailed information about the distribution of environmental suitability and accumulated environmental suitability throughout geographical space can be obtained from niche-based species distribution models when human population density is taken into account. Finally, our findings may serve to subsidize better estimates of areas vulnerable to Chagas disease, as well as to highlight that environmental variables related to human populations should be progressively explored in upcoming macroecological and biogeographical research on vector ecology.

## Acknowledgment

We thank members of the PIBi Lab for productive criticisms and kind comments that helped to improve this manuscript.

## References

[1] Lozano R, Naghavi M, Foreman K, Lim S, Shibuya K, Aboyans V, et al. Global and regional mortality from 235 causes of death for 20 age groups in 1990 and 2010: a systematic analysis for the Global Burden of Disease Study 2010. Lancet 2012;380:2095–128. doi:10.1016/S0140-6736(12)61728-0.

[2] Hotez PJ, Alvarado M, Basáñez M-G, Bolliger I, Bourne R, Boussinesq M, et al. The Global Burden of Disease Study 2010: Interpretation and Implications for the Neglected Tropical Diseases. PLoS Negl Trop Dis 2014;8:e2865. doi:10.1371/journal.pntd.0002865.

[3] Murray KA, Preston N, Allen T, Zambrana-Torrelio C, Hosseini PR, Daszak P. Global biogeography of human infectious diseases. Proc Natl Acad Sci 2015;112:12746–51. doi:10.1073/pnas.1507442112.

[4] Stephens PR, Altizer S, Smith KF, Alonso Aguirre A, Brown JH, Budischak SA, et al. The macroecology of infectious diseases: a new perspective on global-scale drivers of pathogen distributions and impacts. Ecol Lett 2016;19:1159–71. doi:10.1111/ele.12644.

[5] WHO. Investing to overcome the global impact of neglected tropical diseases: third WHO report on neglected diseases 2015. Geneva: 2015. doi:ISBN 978 92 4 156486 1.

[6] Savioli DD and L. Accelerating Work to Overcome the Global Impact of Neglected Tropical Diseases: A Roadmap for Implementation. Geneva: 2012.

[7] Coura JR, Borges-Pereira J. Chagas disease: 100 years after its discovery. A systemic review. Acta Trop 2010;115:5–13. doi:10.1016/j.actatropica.2010.03.008.

[8] Secretaria de Vigilância em Saúde (Ministério da Saúde do Brasil). Doença de Chagas aguda no Brasil: série histórica de 2000 a 2013. Bol Epidemiol 2015;46:1–9. doi:10.1590/S1415-790X2004000400010.

[9] Filho AAF, Sousa AS de, Filho DC, Jansen AM, Andrade GMQ, Britto CFDP de C, et al. II Consenso Brasileiro em Doença de Chagas, 2015. Epidemiol e Serviços Saúde 2016;25:1–10. doi:10.5123/S1679-49742016000500002.

[10] Pereira JM, Almeida PS de, Sousa AV de, Paula AM de, Machado RB, Gurgel-Gonçalves R. Climatic factors influencing triatomine occurence in Central-West Brazil. Mem Inst Oswaldo Cruz 2013;108:335–41. doi:10.1590/S0074-02762013000300012.

[11] Dias JVL, Queiroz DRM, Martins HR, Gorla DE, Pires HHR, Diotaiuti L. Spatial distribution of triatomines in domiciles of an urban area of the Brazilian Southeast Region. Mem Inst Oswaldo Cruz 2016;111:43–50. doi:10.1590/0074-02760150352.

[12] Bezerra CM, Cavalcanti LP de G, Souza R de CM de, Barbosa SE, Xavier SC das C, Jansen AM, et al. Domestic, peridomestic and wild hosts in the transmission of Trypanosoma cruzi in the Caatinga area colonised by Triatoma brasiliensis. Mem Inst Oswaldo Cruz 2014;109:887–98. doi:10.1590/0074-0276140048.

[13] Waleckx E, Gourbière S, Dumonteil E. Intrusive versus domiciliated triatomines and the challenge of adapting vector control practices against Chagas disease. Mem Inst Oswaldo Cruz 2015;110:324–38. doi:10.1590/0074-02760140409.

[14] Costa J, Lorenzo M. Biology, diversity and strategies for the monitoring and control of triatomines - Chagas disease vectors. Mem Inst Oswaldo Cruz 2009;104:46–51. doi:10.1590/S0074-02762009000900008.

[15] Diotaiuti L, Loiola CF, Falcão PL, Dias JCP. The ecology of Triatoma sordida in natural environments in two different regions of the state of Minas Gerais, Brazil. Rev Inst Med Trop Sao Paulo 1993;35:237–45. doi:10.1590/S0036-46651993000300004.

[16] Coura JR, Junqueira ACV, Boia MN, Fernandes O. Chagas disease: from bush to huts and houses. Is it the case of the Brazilian amazon? Mem Inst Oswaldo Cruz 1999;94:379–84. doi:10.1590/S0074-02761999000700074.

[17] Diniz-Filho JAF, Ceccarelli S, Hasperué W, Rabinovich J. Geographical patterns of Triatominae (Heteroptera: Reduviidae) richness and distribution in the Western Hemisphere. Insect Conserv Divers 2013;6:704–14. doi:10.1111/icad.12025.

[18] Costa J, Dornak L, Almeida C, Peterson A. Distributional potential of the Triatoma brasiliensis species complex at present and under scenarios of future climate conditions. Parasit Vectors 2014;7:238. doi:10.1186/1756-3305-7-238.

[19] Ceccarelli S, Rabinovich JE. Global Climate Change Effects on Venezuela’s Vulnerability to Chagas Disease is Linked to the Geographic Distribution of Five Triatomine Species. J Med Entomol 2015;52:1333–43. doi:10.1093/jme/tjv119.

[20] de la Vega GJ, Medone P, Ceccarelli S, Rabinovich J, Schilman PE. Geographical distribution, climatic variability and thermo-tolerance of Chagas disease vectors. Ecography (Cop) 2015;38:851–60. doi:10.1111/ecog.01028.

[21] Parra-Henao G, Suárez-Escudero LC, González-Caro S. Potential Distribution of Chagas Disease Vectors (Hemiptera, Reduviidae, Triatominae) in Colombia, Based on Ecological Niche Modeling. J Trop Med 2016;2016:1–10. doi:10.1155/2016/1439090.

[22] Bustamante DM, Monroy MC, Rodas AG, Juarez JA, Malone JB. Environmental determinants of the distribution of Chagas disease vectors in south-eastern Guatemala. Geospat Health 2007;1:199–211. doi:10.4081/gh.2007.268.

[23] Parra-Henao G, Cardona ÁS, Jaramillo-O N, Quirós-Gómez O. Environmental Determinants of the Distribution of Chagas Disease Vector Triatoma dimidiata in Colombia. Am J Trop Med Hyg 2016;94:767–74. doi:10.4269/ajtmh.15-0197.

[24] Hay SI, Battle KE, Pigott DM, Smith DL, Moyes CL, Bhatt S, et al. Global mapping of infectious disease. Philos Trans R Soc 2013;368:20120250. doi:http://dx.doi.org/10.1098/rstb.2012.0250.

[25] Elith J, Phillips SJ, Hastie T, Dudík M, Chee YE, Yates CJ. A statistical explanation of MaxEnt for ecologists. Divers Distrib 2011;17:43–57. doi:10.1111/j.1472-4642.2010.00725.x.

[26] Austin M. Species distribution models and ecological theory: a critical assessment and some possible new approaches. Ecol Modell 2007;200:1–19.

[27] Peterson AT, Soberón J. Species distribution modeling and ecological niche modeling: getting the concepts right. Nat Conserv 2012;10:102–7.

[28] Peterson a T, Soberón J, Pearson RG, Anderson RP, Martínez-Meyer E, Nakamura M, et al. Ecological niches and geographic distributions. Choice Rev Online 2012;49:49–6266-49–6266. doi:10.5860/CHOICE.49-6266.

[29] Pollock LJ, Tingley R, Morris WK, Golding N, O’Hara RB, Parris KM, et al. Understanding co-occurrence by modelling species simultaneously with a Joint Species Distribution Model (JSDM). Methods Ecol Evol 2014;5:397–406. doi:10.1111/2041-210X.12180.

[30] Anderson RP. When and how should biotic interactions be considered in models of species niches and distributions? J Biogeogr 2017;44:8–17. doi:10.1111/jbi.12825.

[31] Schweiger O, Settele J, Kudrna O, Klotz S, K??hn I. Climate change can cause spatial mismatch of trophically interacting species. Ecology 2008;89:3472–9. doi:10.1890/07-1748.1.

[32] Wisz MS, Pottier J, Kissling WD, Pellissier L, Lenoir J, Damgaard CF, et al. The role of biotic interactions in shaping distributions and realised assemblages of species: Implications for species distribution modelling. Biol Rev 2013;88:15–30. doi:10.1111/j.1469-185X.2012.00235.x.

[33] Heikkinen RK, Luoto M, Virkkala R, Pearson RG, Körber JH. Biotic interactions improve prediction of boreal bird distributions at macro-scales. Glob Ecol Biogeogr 2007;16:754–63. doi:10.1111/j.1466-8238.2007.00345.x.

[34] Araújo MB, Luoto M. The importance of biotic interactions for modelling species distributions under climate change. Glob Ecol Biogeogr 2007;16:743–53. doi:10.1111/j.1466-8238.2007.00359.x.

[35] Araújo MB, Guisan A. Five (or so) challenges for species distribution modelling. J Biogeogr 2006;33:1677–88. doi:10.1111/j.1365-2699.2006.01584.x.

[36] van Proosdij ASJ, Sosef MSM, Wieringa JJ, Raes N. Minimum required number of specimen records to develop accurate species distribution models. Ecography (Cop) 2016;39:542–52. doi:10.1111/ecog.01509.

[37] Hijmans RJ, Cameron SE, Parra JL, Jones PG, Jarvis A. Very high resolution interpolated climate surfaces for global land areas. Int J Climatol 2005;25:1965–78. doi:10.1002/joc.1276.

[38] Center for International Earth Science Information Network - CIESIN - Columbia University. Gridded Population of the World, Version 4 (GPWv4): Population Density, Revision 10. Palisades, NY: NASA Socioeconomic Data and Applications Center (SEDAC) 2017. doi:10.7927/H4DZ068D.

[39] Phillips SJ, Dudík M, Schapire RE. A maximum entropy approach to species distribution modeling. Proc. twenty-first Int. Conf. Mach. Learn., ACM; 2004, p. 83.

[40] Phillips S, Anderson R, Schapire R. Maximum entropy modeling of species geographic distributions. Ecol Modell 2006;190:231–59. doi:10.1016/j.ecolmodel.2005.03.026.

[41] Hutchinson GE. An introduction to population ecology 1978.

[42] Phillips SJ, Dudík M, Elith J, Graham CH, Lehmann A, Leathwick J, et al. Sample selection bias and presence-only distribution models: implications for background and pseudo-absence data. Ecol Appl 2009;19:181–97. doi:10.1890/07-2153.1.

[43] Kramer-Schadt S, Niedballa J, Pilgrim JD, Schröder B, Lindenborn J, Reinfelder V, et al. The importance of correcting for sampling bias in MaxEnt species distribution models. Divers Distrib 2013;19:1366–79. doi:10.1111/ddi.12096.

[44] Fourcade Y, Engler JO, Rödder D, Secondi J. Mapping species distributions with MAXENT using a geographically biased sample of presence data: A performance assessment of methods for correcting sampling bias. PLoS One 2014;9:e97122. doi:10.1371/journal.pone.0097122.

[45] Peterson AT, Papeş M, Soberón J. Rethinking receiver operating characteristic analysis applications in ecological niche modeling. Ecol Modell 2008;213:63–72. doi:10.1016/j.ecolmodel.2007.11.008.

[46] Fourcade Y, Besnard AG, Secondi J. Paintings predict the distribution of species, or the challenge of selecting environmental predictors and evaluation statistics. Glob Ecol Biogeogr 2017;27:245–56. doi:10.1111/geb.12684.

[47] Lobo JM, Jiménez-Valverde A, Real R. AUC: a misleading measure of the performance of predictive distribution models. Glob Ecol Biogeogr 2008;17:145–51.

[48] Jiménez-Valverde A. Insights into the area under the receiver operating characteristic curve (AUC) as a discrimination measure in species distribution modelling. Glob Ecol Biogeogr 2011:no-no.

[49] Moreno-Amat E, Mateo RG, Nieto-Lugilde D, Morueta-Holme N, Svenning JC, García-Amorena I. Impact of model complexity on cross-temporal transferability in Maxent species distribution models: An assessment using paleobotanical data. Ecol Modell 2015;312:308–17. doi:10.1016/j.ecolmodel.2015.05.035.

[50] Anderson RP, Gonzalez I. Species-specific tuning increases robustness to sampling bias in models of species distributions: An implementation with Maxent. Ecol Modell 2011;222:2796–811. doi:10.1016/j.ecolmodel.2011.04.011.

[51] Poo-Muñoz DA, Escobar LE, Peterson AT, Astorga F, Organ JF, Medina-Vogel G. Galictis cuja (Mammalia): an update of current knowledge and geographic distribution. Iheringia Série Zool 2014;104:341–6. doi:10.1590/1678-476620141043341346.

[52] Rangel TF, Diniz-Filho JAF, Bini LM. SAM: a comprehensive application for Spatial Analysis in Macroecology. Ecography (Cop) 2010;33:46–50. doi:10.1111/j.1600-0587.2009.06299.x.

[53] Medone P, Ceccarelli S, Parham PE, Figuera A, Rabinovich JE. The impact of climate change on the geographical distribution of two vectors of Chagas disease: implications for the force of infection. Philos Trans R Soc B Biol Sci 2015;370:20130560– 20130560. doi:10.1098/rstb.2013.0560.

[54] Cajo DJY, Moreno M, Chaguamate L, Valencia N, Ayala VR. Aplicación de Modelos de Nicho Ecológico para estudios Epidemiológicos: Triatoma dimidiata, vector de la 1 Enfermedad de Chagas en Ecuador. Rev Politécnica 2016;37:88–92.

[55] de Melo França L, Fortier DC, Bocchiglieri A, Dantas MAT, Liparini A, Cherkinsky A, et al. Radiocarbon dating and stable isotopes analyses of Caiman latirostris (Daudin, 1801)(Crocodylia, Alligatoridae) from the late Pleistocene of Northeastern Brazil, with comments on spatial distribution of the species. Quat Int 2014;352:159–63.

[56] Waleckx E, Salas R, Huamán N, Buitrago R, Bosseno MF, Aliaga C, et al. New insights on the Chagas disease main vector Triatoma infestans (Reduviidae, Triatominae) brought by the genetic analysis of Bolivian sylvatic populations. Infect Genet Evol 2011;11:1045–57. doi:10.1016/j.meegid.2011.03.020.

[57] Schofield CJ. Triatominae: biology and control. Bognor Regis, UK: Eurocommunica Publications; 1994.

[58] Galvão C. Vetores da Doença de Chagas no Brasil. Curitiba: SciELO Books - Sociedade Brasileira de Zoologia; 2014.

[59] Altizer S, Ostfeld RS, Johnson PTJ, Kutz S, Harvell CD. Climate Change and Infectious Diseases: From Evidence to a Predictive Framework. Science (80-) 2013;341:514–9. doi:10.1126/science.1239401.

[60] Jones KE, Patel NG, Levy MA, Storeygard A, Balk D, Gittleman JL, et al. Global trends in emerging infectious diseases. Nature 2008;451:990–3. doi:10.1038/nature06536.

[61] Smith KF, Goldberg M, Rosenthal S, Carlson L, Chen J, Chen C, et al. Global rise in human infectious disease outbreaks. J R Soc Interface 2014;11:20140950–20140950. doi:10.1098/rsif.2014.0950.

